# Wild-type and single-O-antigen repeat outer-membrane vesicles induce equivalent protection against homologous and heterologous *Salmonella* challenge

**DOI:** 10.1101/2024.07.03.601871

**Authors:** Areej Alshayea, Sian Emily Jossi, Edith Marcial-Juárez, Marisol Pérez-Toledo, Ruby Persaud, Anna Elizabeth Schager, Daniel Nyberg Larsen, Gvantsa Gutishvili, Jamie Pillaye, Fernanda Escobar-Riquelme, Kubra Aksu, Jack A. Bryant, William G. Horsnell, Manuel Banzhaf, Jakub Zbigniew Kaczmarek, Peter Højrup, James C. Gumbart, Ian Robert Henderson, Vassiliy N. Bavro, Constantino López-Macías, Adam F. Cunningham

## Abstract

Lipopolysaccharide O-antigen is an immunodominant target of protective antibodies. Variation in O-antigen structures limits antibody-mediated cross-protection between closely-related pathogens including *Salmonella* Typhimurium (STm) and *S*. Enteritidis (SEn). Bacterial outer membrane vesicles (OMV) are vaccine platforms presenting surface antigens in their natural conformations. To assess how O-antigen lengths impact antibody responses and control of homologous or heterologous infection, mice were immunized with STm-OMV containing wild-type O-antigen unit repeats (wt-OMV), ≤1 O-antigen unit (wzy-OMV), or no O-antigen units (wbaP-OMV) respectively and challenged with either STm or SEn. Unexpectedly, anti-STm LPS IgG and protection to STm were comparable after immunization with either wt-OMV or wzy-OMV. Anti-porin responses were elevated after immunization with wzy-OMV and wbaP-OMV. A single immunization with any OMV induced minimal cross-protection against SEn, except in blood. In contrast, boosting with O-antigen-expressing OMV enhanced control of SEn infections by >10-fold. These results suggest that i) Antibody to single or variable-length O-antigen units are comparably protective against *Salmonella*; ii) Antigens other than immunodominant O-antigens may be targets of cross-reactive antibodies that moderate bacterial burdens; iii) Boosting can enhance the level of cross-protection against related *Salmonella* serovars and iv) High tissue burdens of *Salmonella* can be present in the absence of detectable bacteraemia.

## Introduction

Antibodies induced to vaccines and following natural infection play an important role in protecting from viral and bacterial pathogens. Generating protective antibody responses requires the selection of B cells that recognize antigens that are presented to the immune system in the same conformation as produced by the bacterial pathogen. The germinal centre (GC) pathway plays a critical role in this selection process and GC are essential for the induction of maximal antibody-responses, as well as for the generation of B cell memory and long-lived, high-affinity plasma cells, which reside in sites like the spleen and bone marrow ^1, 2, 3^. Although antibody levels typically wane after vaccination or infection, this does not necessarily result in an equivalent decrease in the functionality of antibody responses, which can be maintained, or even enhanced, over time ^4^. Protection against re-infection with Gram-negative bacteria is associated with antibodies to the bacterial cell-surface and all licensed human vaccines contain cell-surface or secreted antigens. In the absence of a capsular polysaccharide, the immunodominant O-antigen polysaccharide sub-component of lipopolysaccharide (LPS) is a key target of antibodies that provide serovar-specific protection against infection, but limited cross-protection against serovars that express a different O-antigen ^5, 6, 7, 8, 9, 10, 11, 12, 13,14^. An elegant example can be seen after infection and challenge with different *Salmonella* serovars and isogenic strains that differ in their LPS O-antigen expression ^6^. In these studies Hormaeche and colleagues showed that there is limited cross-protection between *Salmonella* Typhimurium (STm), expressing the O4 antigen, and S. Enteritidis (SEn), expressing O9 ^6, 15^,despite these O-antigen variants from STm and SEn sharing the same trisaccharide carbohydrate backbone ^15^. This critical potential protective value of antibodies to O-antigen results in the O-antigen being included in many vaccines in development against Gram-negative bacteria, including non-typhoidal *Salmonella* infections ^16^. While antibodies to O-antigen are clearly important, antibodies to non-LPS antigens can also contribute to protection, although which other antigens are protective is less well explored or understood. Examples of *Salmonella* antigens that may be protective include the trimeric porin proteins and the trimeric autotransporter SadA from STm ^4, 17, 18, 19, 20^. Indeed, human subunit vaccines, such as vaccines against pertussis or meningococcal B infections do not contain LPS while providing effective protection against disease ^21, 22, 23^. Therefore, the combination of appropriate antigen targeting and the nature of the immune response to that antigen combine to influence overall efficacy of antibody responses.

Since antibodies to both LPS O-antigen and non-LPS antigens can be protective, the inclusion of multiple antigens within a vaccine may be desirable. However, generating multi-antigen-containing subunit vaccines can increase vaccine cost and affordability, as each individual antigen needs to be synthesised and purified. Outer-membrane vesicles (OMV), are membrane blebs released naturally from bacteria, and the rate of bacterial shedding of OMV from Gram-negative bacteria can be enhanced through the genetic deletion of *tolR* ^24^. OMV hold promise as vaccine vectors not least because they contain cell envelope antigens, including LPS and surface proteins, such as porins, in their native conformation ^25, 26^. Nevertheless, how to best exploit the OMV platform remains unclear. For example, although O-antigen is a target of protective antibodies, O-antigen chains can also restrict antibody access to non-O-antigen targets on the bacterial surface ^7, 27, 28^, reducing the immune response to these antigens. Thus, limiting the amount of O-antigen in OMV may enhance antibody responses to non-LPS antigens after immunization. Bacterial *wbaP* or *wzy* knock-out mutants produce LPS that contain the lipid A and core oligosaccharide components, but either lack O-antigen repeats altogether, or express an LPS molecule containing a maximum of one O-antigen repeat respectively ^29, 30^(Fig. 1A). Crucially, immunizing with OMV derived from wzy mutants may result in the induction of anti-O-antigen antibodies, and structural biology studies of antibody binding to *Salmonella* O-antigen suggest that a single epitope could be contained within a single O-antigen repeat ^31, 32^. To determine how levels of LPS O-antigen in OMV affect the antibody responses induced, we assessed immune responses to distinct preparations of OMV from different STm mutants (Fig. 1A). Alongside, we evaluated the capacity of immunization with different OMV to control bacterial infection after challenge with STm (which expresses O4-type LPS), and to reduce bacterial burdens after infection with SEn (which expresses O9-type LPS). Our studies demonstrate how O-antigen length impacts the levels of cross-serovar antibodies induced, and how boosting has an unexpected impact on the ability to control infections caused by bacteria expressing different O-antigen types.

**Figure 1.**
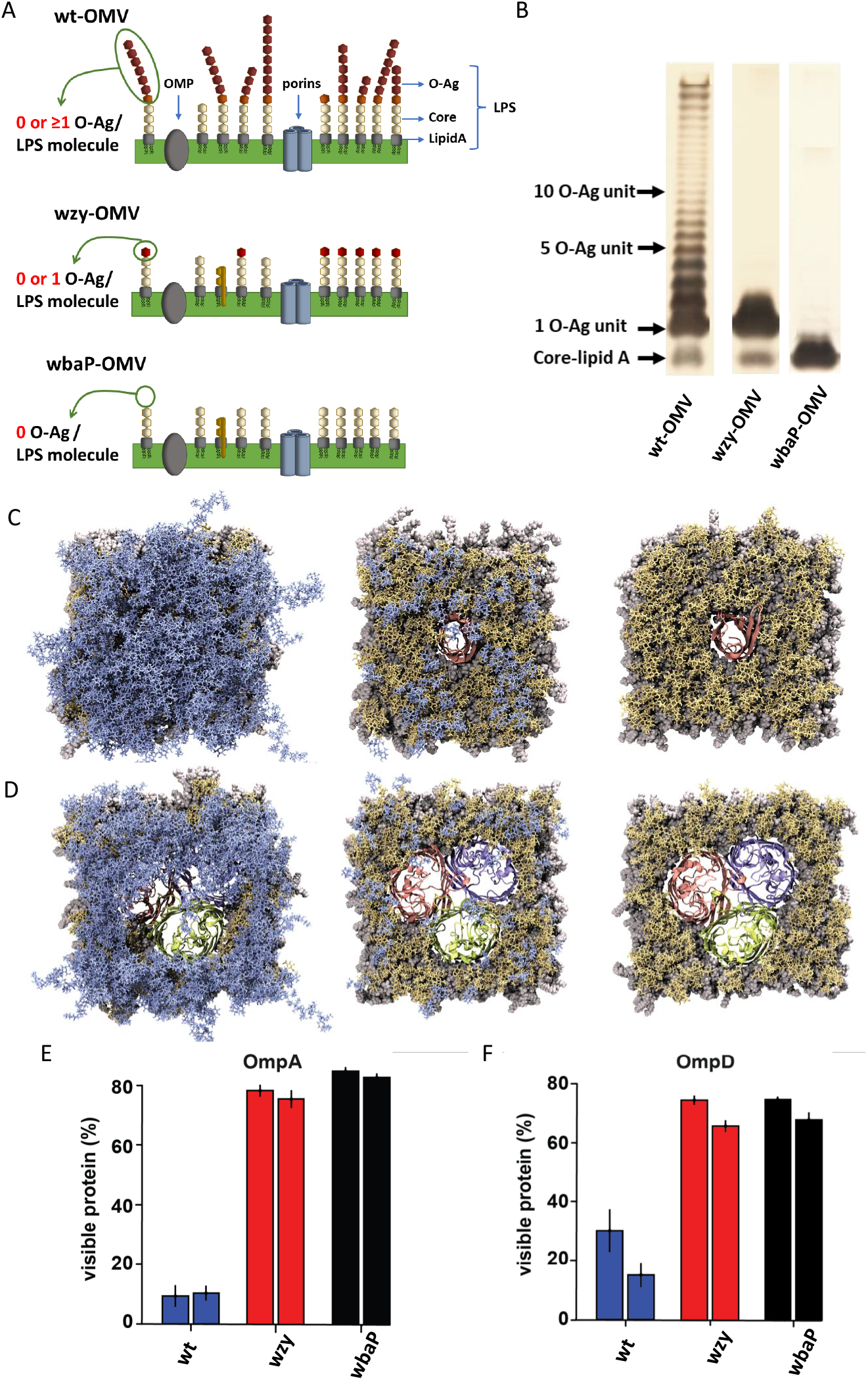
The effect of different O-antigen repeat length in the OMV-producing STm mutants. **(A)** Schematic diagram of the LPS composition in the wild type (wt) and the two different OMV-producing mutants – wzy and wbaP respectively. **(B)** A silver-stain SDS-PAGE confirming the distribution of O-antigen repeats in wt, wzy and wbaP. **(C, D)** Visualisation of the effect of the number of O-antigen repeats on visibility of the monomeric porin OmpA **(C)** and trimeric porin OmpD **(D)** respectively. Results from molecular dynamics simulations illustrating (from right to left) discrete STm wt, wzy and wbaP mutants containing 6-11 or 0-1 or 0 O-antigen units (in blue) respectively; view from outside of the cell perpendicular to the membrane plane. (**E, F**) Bar plots show the percentage of OmpA (**E**) and OmpD (**F**) protein visible for each of the LPS types, with two columns corresponding to the two different replicas of simulations for the same model.

## Materials and Methods

### Bacterial strains, antigens and mice

*Salmonella enterica* serovars Typhimurium (STm) and Enteritidis (SEn) were used for experiments ^7^. STm SL1344 and 14028 are standard laboratory wt strains, D23580 is a wt strain isolated from a child in Malawi ^33^. SEn D24954 is a wt virulent strain isolated from a child in Malawi ^34^. STm 14028 Δ*tolR*, 14028 Δ*wzy* and 14028 Δ*wbaP* mutants were obtained from an ordered single gene deletion library ^35^. LPS from STm was obtained from Enzo Life Sciences LTD. This LPS contains O-antigen, core and lipid A components. Purified porins from STm were produced from STm 14028 as previously described ^7, 17^.

CD1 female wt mice, aged 6-8 weeks were purchased from Envigo RMS (UK). All mice used throughout this study were bred and maintained under specific-pathogen free conditions at the Biomedical Service Unit (BMSU) at University of Birmingham. All animal experiments were performed in accordance with UK Home Office regulations (project licence P06779746).

### Generation of OMV producing strains and the purification of OMV

Enhanced OMV producing strains were generated by disruption of *tolR* in STm 14028, STm 14028 Δ*wzy*, and STm 14028 Δ*wbaP* by insertion of a kanamycin resistance cassette (kan) using standard methods ^36^. Gene deletion was confirmed by PCR and LPS O-antigen was detected by silver staining purified OMV. Purification of OMV was performed as described elsewhere ^8^. Briefly, 300 ml cultures of each Δ*tolR* bacterial strain was grown to OD600 of 1 and bacteria harvested by centrifugation (4000 x g, 10 minutes, 4°C). Supernatants were filtered using a 0.22 µm 500 ml filter unit before filtrates were spun in an ultra-centrifuge for 2 hours (4°C, 186,000 x g). Pellets were washed with cold sterile PBS and centrifuged again for 1 hour with the same conditions. Finally, pellets were resuspended in cold PBS and filtered again using a small 0.22 µm filter. The protein concentration of OMV was determined by BCA assay and OMV were examined by SDS-PAGE analysis. Samples were prepared by heating OMV for 10 minutes at 70 °C with NuPAGE LDS sample buffer and sample reducing agent (Invitrogen). Samples were loaded to a NuPAGE 4-12% gradient Bis-Tris gel (Invitrogen) and stained with Coomassie Blue stain (Thermo Fisher Scientific) or with a Pierce Silver Stain Kit (Thermo Scientific) following the manufacturer’s instructions. Preparations of OMV were stored at 4°C.

### Sample Preparation for mass spectrometry analysis

Each OMV sample was reduced by addition of dithiothreitol (DTT) to a final concentration of 10 mM, followed by incubation at 57°C for 30 min. Alkylation was performed by addition of iodoacetamide (IAA) to a final conc. of 24 mM and incubated for 20 min in the dark. The alkylation’s was quenched by adding 1μl 0.05 M DTT to a 60µl sample. OMV samples were then digested overnight at at 37 °C with 2% w/w in-house methylated trypsin ^37^. Each OMV sample was then micro purified essentially as described by Rappsilber et al. ^38^, followed by lyophilization and resuspension in 0.1% formic acid (FA).

The mass spectrometric analysis was performed on a Thermo Scientific Orbitrap Exploris 480™ coupled to an EASY-nLC1000 Liquid Chromatography system (Thermo Fisher Scientific), utilizing a 3 cm trap column (100 μm inner diameter, 5 μm Reprosilpur 120 C18, Dr. Maisch GmbH, Germany) and an 18 cm analytical column (75 μm inner diameter, 3 μm Reprosilpur 120 C18). The method utilized was a standard top 10, 60 min gradient using 120,000 MS1 and 30,000 MS2 resolution. The data analysis was performed using the Proteome Discoverer Software version 2.5 (Thermo Fisher Scientific).

### Immunization and infection of mice

Mice were immunised intraperitonially (i.p.) with 1 μg STm-OMV in PBS, where mass is based on protein content, for the times described in the results. Mice were challenged i.p. with 1×10^4^ CFU STm D23580 or 1×10^4^ SEn D24954 in 200 μL sterile PBS. At the end of experiments, blood was collected through cardiac puncture. Blood cultures were performed immediately after isolation of blood by plating 100 μl of blood onto an LB agar plate. Sera were obtained from bloods allowed to stand for 1 hour. Bacterial numbers in the spleens and livers from infected mice were assessed by direct culture of organs processed by passing through a 70 μm cell strainer.

### Preparation of cells for ELISPOT and flow cytometry

Cell suspensions were prepared from spleens and bone marrows for use in ELISPOT and flow cytometry assays using previously described methods ^8^. Splenocytes were isolated by mashing spleens through a 70 μm cell-strainer and bone marrow cells were isolated by flushing both the tibia and femur of one hind leg with R-10 media and erythrocytes lysed using standard protocols.

### B cell Enzyme-Linked ImmunoSpot (ELISPOT)

Antigen-specific antibody secreting cells (ASC) in the spleen and bone marrow were determined by ELISPOT using previously described methods ^8, 39^. MultiScreen 96-well filtration plates (Merck Millipore) were coated with LPS or porins (5 µg/ml) in PBS overnight at 4 °C. Wells were washed with PBS and blocked with R-10 media for 1 hour at 37 °C. 5 × 10^5^ cells per well were added and plates incubated at 37 °C with 5% CO2 for 6 hours before washing with PBS with 0.05% Tween 20. Alkaline phosphatase conjugated anti-IgG and anti-IgM (1:1000; Southern Biotech) diluted in PBS was added overnight at 4 °C. Signal was detected by adding SIGMAFAST 5-Bromo-4-chloro-3-indolyl phosphate / Nitro blue tetrazolium (BCIP/NBT) tablets according to the manufacturer’s instructions. Spots were counted using an AID ELISPOT plate reader and AID software version 7.0 (AID GmbH). ASC counts were presented as spot-forming units (SFUs) per 1 × 10^6^ cells.

### Flow cytometry

Flow cytometry was performed as described previously ^8, 39, 40^. Splenocytes were resuspended in FACS buffer in a 96-well V-bottom plates at 2.5 × 10^6^ cells with blocking reagent (anti-CD16/CD32) for 15 minutes at 4 °C. After blocking, the antibodies in FACS buffer were added and incubated on ice for 25 minutes in the dark. Antibodies used are described in Supplementary Table 1. After washing, viability dye was added and cells incubated in PBS for 15 minutes at room temperature in the dark. Cells were then fixed with 0.1% w/v paraformaldehyde (PFA) or permeabilised for intracellular staining using a Foxp3 Fixation/Permeabilization kit (eBioscience) before staining. Samples were acquired using a BD LSRII Fortessa flow cytometer and analysed using FlowJo 10.1 software (TreeStar).

### Enzyme-linked immunosorbent assay (ELISA)

ELISAs were performed as previously described ^7, 17^. Maxisorp 96 well NUNC plates (Thermo Fisher Scientific) were coated overnight at 4 °C with 5 μg/ml of antigen or bacteria diluted in carbonate coating buffer. Plates were blocked with 1% BSA (Sigma-Aldrich) in PBS and incubated for 1 hour at 37°C. Sera were serially diluted in 3-fold steps and incubated for 1 hour at 37°C. After washing, AP-conjugated goat anti-mouse immunoglobulin antibodies were added to detect IgM (1 in 2000) or IgG (1 in 1000) (Southern Biotech). Signal was detected with Sigma FAST P-nitrophenyl phosphate (pNPP) tablets (Sigma-Aldrich) dissolved in water according to manufacturer instructions. Plates were read by measuring the absorbance of each well at OD405 nm on a SpectraMax ABS plus plate reader (Molecular Devices) using SoftMax pro software version 6.5. Titres were calculated by identifying the dilution at which the signal reached a set OD405.

### Structural modelling and analysis

We constructed multiple atomistic models and carried out molecular dynamics simulations to investigate the dynamical behaviour of the outer membrane and embedded porins. A serovar-specific OmpD trimer was modelled based on a homology model used previously ^7^. The model of OmpA was taken from AlphaFold2 (P02936), and the signal sequence (residues 1-21) was removed ^41^. Each protein was placed into five distinct outer-membrane models using CHARMM-GUI ^42^. In all the membranes, the inner leaflet consisted of 75% POPE and 25% POPG lipids ^43^. The outer leaflet consisted of 100% LPS molecules, featuring identical Lipid A and core oligosaccharide units but varying in the length and composition of O-antigen chains. O4 antigen was modelled representing the dominant LPS form expressed by STm, with three different lengths of repeating units. Our three membranes included (1) wbaP, representing the outer leaflet with only Lipid A and core oligosaccharide; (2) wzy, incorporating one additional O-antigen unit; and (3) wt, containing a random number of repeating O-antigen units, ranging from 6 to 11, for each LPS molecule. To neutralize the large negative charge on the LPS, we introduced Mg^2+^ and Ca^2+^ ions, followed by the addition of Na^+^ and Cl^−^ ions to water to achieve a salt concentration of 0.15 M. System sizes ranged from 100-300,000 atoms, depending on the protein and length of the O-antigen chains.

Simulations were conducted using NAMD3 ^44^ for 6 different models (OmpA and OmpD trimer, each in one of three STm model outer membranes), with two replicas for each system (Fig. 1C, D). Simulations were run at a constant temperature of 310 K and constant pressure of 1 atm (separately maintained in the membrane plane and normal to the membrane). The CHARMM36m force field for proteins ^45^ and CHARMM36 force for lipids ^46^ were used. Initially, we employed a 2-fs time step, allowing the system to equilibrate for 40 ns (not included in the analysis). Subsequently, we implemented hydrogen mass repartitioning (HMR) ^47^ and used a uniform 4-fs time step to simulate an additional 1.5 μs for each system (18 μs in total). For analysis and visualization, we employed Visual Molecular Dynamics (VMD) ^48^. Analysis in Fig. 1E, F were obtained by averaging over the last 0.5 μs, at which point the systems are already equilibrated ^7^.

For the purposes of structural analysis of LPS recognition by antibodies we utilised the experimental structures of the following antibody-polysaccharide complexes: 3BZ4.pdb and 3C6S.pdb showing the decasaccharide and undecasaccharide respectively being bound by an anti-LPS monoclonal antibody ^49^. We also analysed IGG1-lambda SE155-4 FAB binding to both synthetic and natural saccharide epitopes in the following structures: 1MFE.pd ^31^; 1MFA.pdb ^32^; 1MFE.pdb ^50^, alongside the related *Shigella* data from Fab SYA/J6 (1M7D.pdb; 1M7I ^51^; ^52^), all presenting binding to trisaccharide epitopes, as well as the 1M7I ^51^ showing a pentasaccharide recognition.

### Statistical analysis

GraphPad Prism version 8 was used for data analysis. Statistical differences were determined using the Mann-Whitney U test or Kruskal-Wallis with Dunn’s multiple comparisons test. Statistical significance was accepted when p ≤ 0.05

## Results

### O-antigen length in OMV does not affect protein composition but modulates antigen exposure

To generate a panel of STm strains producing OMV, the tolR gene from wild-type STm 14028, and its isogenic Δ*wzy* and Δ*wbaP* mutants, was replaced by allelic exchange with a kanamycin resistance cassette. For clarity, in the text below, the OMV generated from *tolR* mutants are referred to as wild-type (wt)-OMV, as they express LPS O-antigens of variable lengths. Additionally, wzy-OMV or wbaP-OMV refer to OMV purified from bacterial cultures of *tolR wzy* or *tolR wbaP* double-mutants respectively (Fig. 1A), which express LPS containing either a single or no O-antigen repeat, as confirmed by silver stain SDS-PAGE (Fig. 1B).

All strains produced OMV with similar protein profiles (Supplementary Fig. 1). Moreover, assessment of the OMV by mass spectrometry revealed a high level of conservation of proteins between different OMV, with OmpA and the trimeric porins included among the more commonly detected proteins. Over 65% of the proteins identified were found in all three mutants, while 78% were found in two out of three mutants, with the majority of differences accounted by bystander cytoplasmic proteins copurifying with the OMV. This suggests that despite the differences in O-antigen expression between OMV, they retain a similar profile of cell-envelope proteins within them.

Based on our previous data ^7^, we anticipated that there may be a significant difference in visibility of proteinaceous antigens in those mutants, with wbaP allowing for higher porin accessibility. To assess this in more detail, we modelled the LPS layer in the OMV surrounding the monomeric porin OmpA (Fig. 1C, E) and the representative trimeric porin OmpD respectively (Fig. 1D, F) by all-atom molecular dynamics (MD). The results from protein visibility analyses (Fig. 1E, F) confirm the anticipated increase in visibility of monomeric porins for both the wzy and wbaP mutants, while for the wt (Fig. 1C, E), these are fully occluded. In contrast, the trimeric porins, such as OmpD are predicted to be continuously visible in all LPS scenarios, including for IgG binding as previously shown (Fig. 1D, F) ^7^. This suggests that OMV containing truncated O-antigens could be effective vehicles for presentation of proteinaceous antigens.

### wt, wzy and wbaP-OMV induce similar germinal centre responses, but antigen-specific responses vary depending upon O-antigen content

OMV are complex antigens containing protein, carbohydrate, and lipid antigens, and variations in the content of antigens can alter their immunological properties ^53, 54^. To examine the impact of reduced or absent O-antigen levels on the immune response to OMV, mice were immunized with 1 μg of each OMV for 21 or 35 days. Some mice were boosted at 35 days and 21 days later splenic GC B cell and total IgM+ and IgG+ plasma(blast) responses assessed. Immunization with wt, wzy or wbaP-OMV resulted in increases in the median proportions and numbers of GC B cells compared to non-immunized controls (Fig. 2A and gating strategy in Supplementary Fig. 2A). Increased total splenic IgM and IgG ASC proportions and numbers were also detected 21 days after primary immunization but less consistently thereafter (Gating strategy in Supplementary Fig. 2B and data in Supplementary Fig. 3). We next examined whether there were differences in the response to porins and LPS by using ELISPOT (representative ELISPOT shown in Supplementary Fig. 4A) to assess responses in the spleen and bone marrow (BM), key sites for the maintenance of long-lived plasma cell responses ^55^. In both the spleen and the BM, IgM+ (Supplementary Fig. 4B and C) and IgG+ ASC (Fig. 2B and C) were detected against LPS and porins post-immunization with any of the OMV types. Mice immunized with wbaP-OMV typically had lower median frequencies of anti-LPS IgM+ and IgG+ ASC than mice immunized with wt and wzy-OMV. In contrast, wt and wzy-OMV induced similar median anti-LPS ASC responses as each other. Mice immunized with wzy-OMV or wbaP-OMV typically had a higher frequency of anti-porin IgG+ ASC than mice immunised with wt-OMV. In all cases, for each OMV type, the highest median ASC responses against LPS or porins were observed 21 days after boosting.

**Figure 2.**
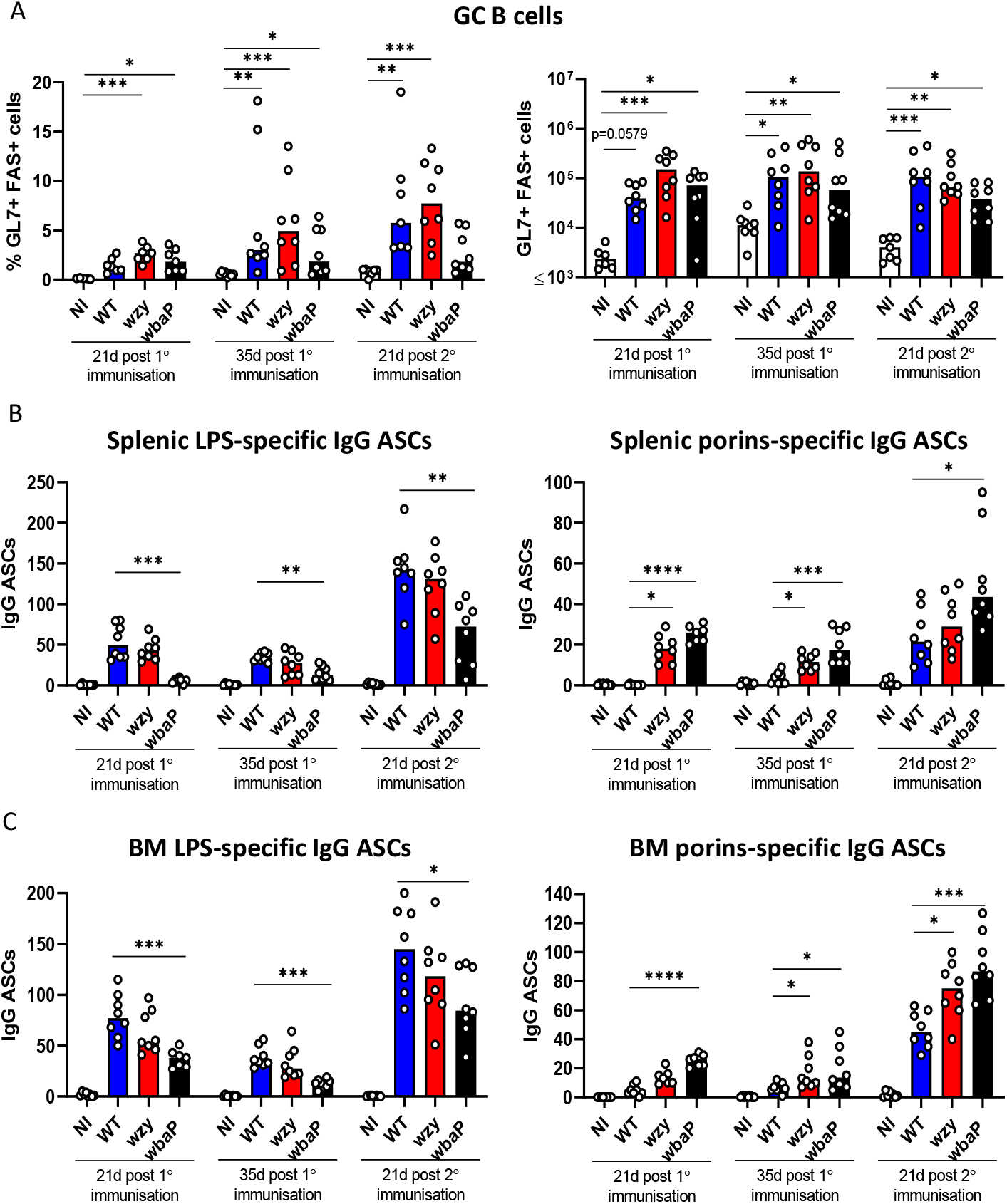
Germinal centre and IgG antibody secreting cells in spleen and bone marrow specific to LPS and porin induced after immunization with OMV. **(A)** Proportion (from total B cells) and the total number of splenic germinal centre B cells (GL7^+^FAS^+^ B cells) in wt mice immunized i.p. with 1μg of OMV for 21 or 35 days or boosted at 35 days for a further 21 days. **(B)** Frequencies of splenic LPS or porin-specific IgG antibody-secreting cells were determined by ELISPOT. **(C)** Frequencies of bone marrow LPS or porin-specific IgG antibody-secreting cells. Bars show medians and individual points represent counts from a single mouse. NI = non-immunized, wt = wt-OMV, wzy = wzy-OMV, wbaP = wbaP-OMV. * = p≤0.05, ** = p≤0.01, *** = p≤0.005, **** = p≤0.001. Graphs show the combined data from 2 independent experiments.

**Figure 3.**
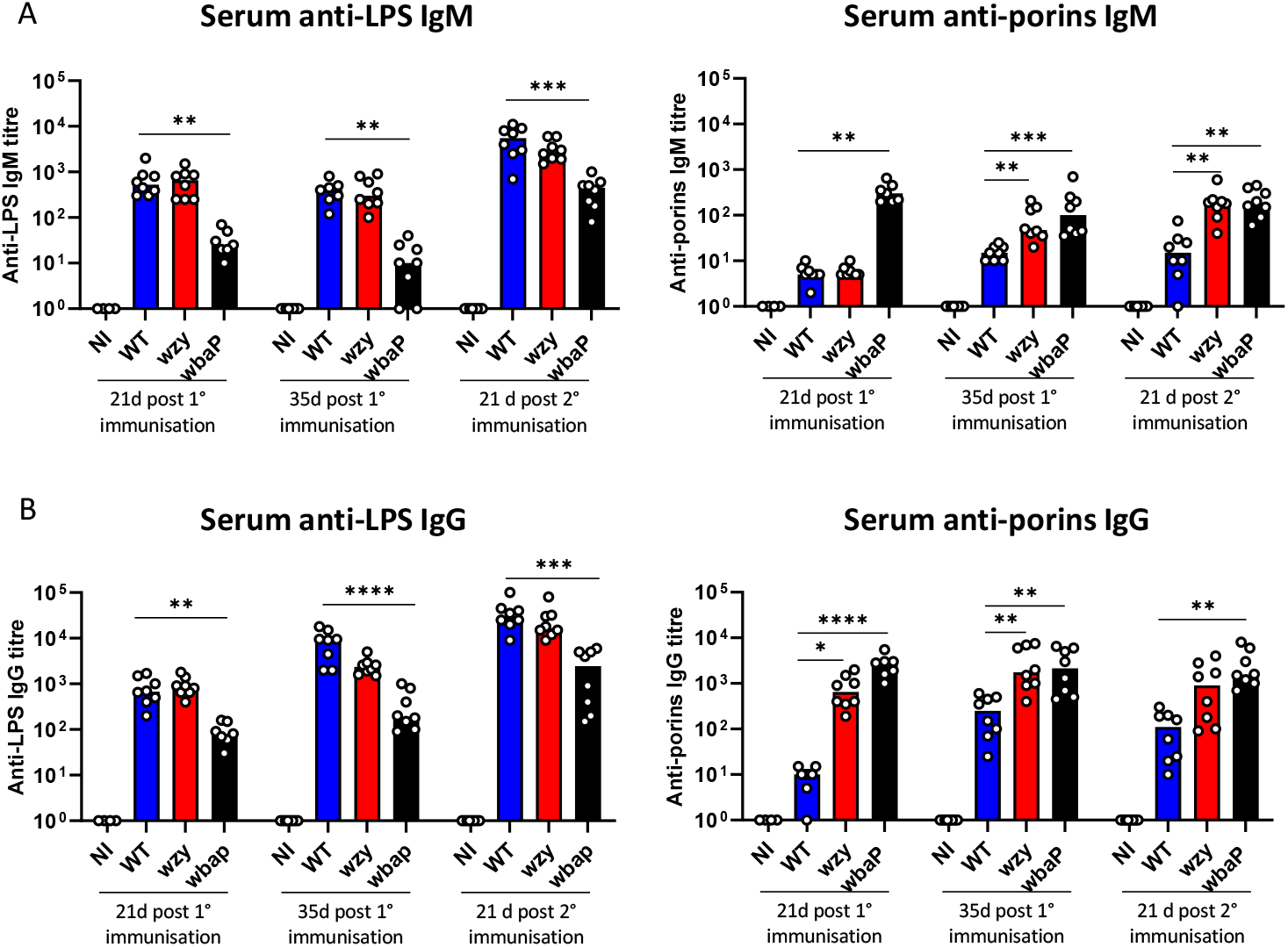
Anti-LPS and anti-porin IgM and IgG serum titres induced to OMV. wt mice were immunized i.p. with 1μg of OMV for 21 or 35 days or boosted at 35 days for a further 21 days. **(A)** Anti-LPS and anti-porin IgM titers determined by ELISA. **(B)** Anti-LPS and anti-porin IgG titers determined by ELISA. Bars show medians and individual points represent counts from a single mouse. NI = non-immunized, wt = wt-OMV, wzy = wzy-OMV, wbaP = wbaP-OMV. * = p≤0.05, ** = p≤0.01, *** = p≤0.005, **** = p≤0.001. Graphs show the results from sera generated from 2 independent experiments.

**Figure 4.**
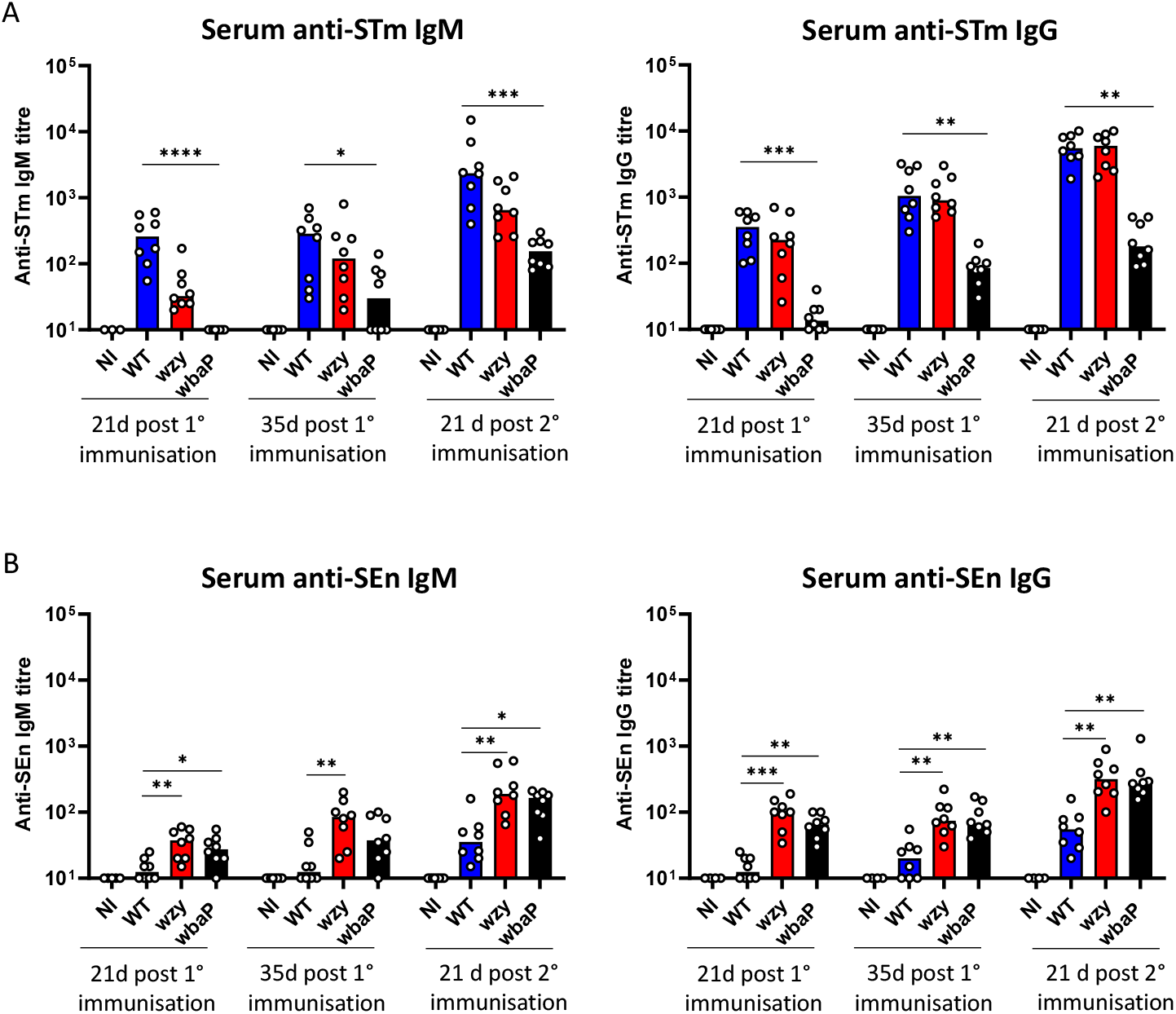
Anti-STm and anti-SEn IgM and IgG serum titres induced to OMV. wt mice were immunized i.p. with 1μg of OMV for 21 or 35 days or boosted at 35 days for a further 21 days. **(A)** Anti-STm IgM and IgG titres determined by ELISA. **(B)** Anti-STm IgM and IgG titres determined by ELISA. Bars show medians and individual points represent a single mouse. NI = non-immunized, wt = wt-OMV, wzy = wzy-OMV, wbaP = wbaP-OMV. * = p≤0.05, ** = p≤0.01, *** = p≤0.005, **** = p≤0.001. Graphs show the results from sera generated from 2 independent experiments.

Assessment of serum antibody responses by ELISA showed that anti-LPS IgM or IgG responses were similar after immunization with either wt or wzy-OMV (Fig. 3A, B) and were higher than after immunization with wbaP-OMV. In contrast to the anti-LPS response, the anti-porin IgM and IgG responses were more similar between the wzy and wbaP-OMV groups than either group to the wt-OMV group. Thus, wzy-OMV induce anti-LPS responses similar to those induced by wt-OMV, but anti-porin responses are more similar to those induced by wbaP-OMV.

### Immunization with wzy-OMV induces similar anti-STm IgG as wt-OMV but enhanced IgG to SEn

Next, we assessed how the serological response to the individual antigens reflected the binding of serum antibodies to whole STm bacteria (Fig. 4 A,B). In addition, we assessed the level of cross-reactivity of these antibodies to SEn (Fig. 4 C, D), which expresses a related, but different, O-antigen. At each comparable time-point, IgM and IgG binding to STm was similar between the wt or wzy-OMV groups (Fig. 4A). Median IgM and IgG titres were lower for sera from the wbaP-OMV immunized groups at each time-point, although binding did increase with time and after boosting. The increase in anti-STm in the wbaP-OMV immunized groups suggested the antibody response against non-O-antigens could evolve over time and be enhanced by boosting (Fig. 4A). To examine this, the same sera generated after immunization with the different OMV were tested against SEn, which expresses O9 O-antigen rather than the O4 O-antigen expressed by STm (Fig. 4C, D). Against SEn, there was little change in IgM and IgG titres between days 21 and 35 after primary immunization, although these anti-SEn median titres were highest after immunization with wzy-OMV or wbaP-OMV. After boosting, anti-SEn titres increased, with the median titres highest in the wzy and wbaP-OMV groups. Thus, immunization with wt and wzy-OMV induce similar levels of anti-STm antibodies, but wzy and wbaP-OMV induce higher levels of cross-reactive antibodies at equivalent times post-immunization.

### Boosting with OMV enhances control of bacterial infection after challenge with SEn

OMV can confer extensive protection against the parent serovar ^8, 56, 57^. Accordingly, mice immunized once for 21 days with the different OMV types and then challenged with STm all had lower bacterial burdens when compared to non-immunized mice, with no bacteria detected in the blood of any of the OMV-immunized mice (Fig. 5A). STm was also not detected in the spleens or livers of most mice immunized with wt-OMV or wzy-OMV STm (Fig. 5A). This was not the case after immunization with wbaP-OMV, where bacteria were detected in the spleens and livers of all mice. This is consistent with O-antigen being a major determinant of antibody-mediated protection. After boosting with wbaP-OMV and challenge with STm no bacteria were detected in the spleens or livers of 4 out of 8 mice (Fig. 5B). Since immunization with wzy and wbaP-OMV induced cross-serovar binding antibodies (Fig. 4), the capacity of immunisation with the different OMV to reduce bacterial burdens following challenge with SEn was assessed (Fig. 5 C, D). In these experiments, any reduction in bacterial numbers cannot be attributable to antibody to O4 O-antigen, as this O-antigen type is not expressed by SEn. Notably, bacteria were not detectable in the blood of most OMV-immunized and SEn-challenged mice (Fig. 5C, D). After SEn challenge of mice immunized once for 21 days with wt, wzy or wbaP-OMV respectively there was a median 8-, 2- and 6-fold reduction in bacterial numbers in their livers and a median 5-, 1.5- and 5-fold reduction in their spleens respectively (Fig. 5C). After challenge of mice boosted with wt, wzy and wbaP-OMV, there was a marked enhancement in the reduction of bacterial burdens in tissues compared to non-immunized controls (Fig. 5D). In the livers of wt, wzy and wbaP-OMV-boosted mice the reduction was 24-, 18- and 7-fold respectively. In the spleens of wt, wzy and wbaP-OMV-boosted mice the reduction was 44-, 58- and 16-fold respectively. Thus, boosting can significantly enhance cross-protection in an O4-antigen-independent manner.

**Figure 5.**
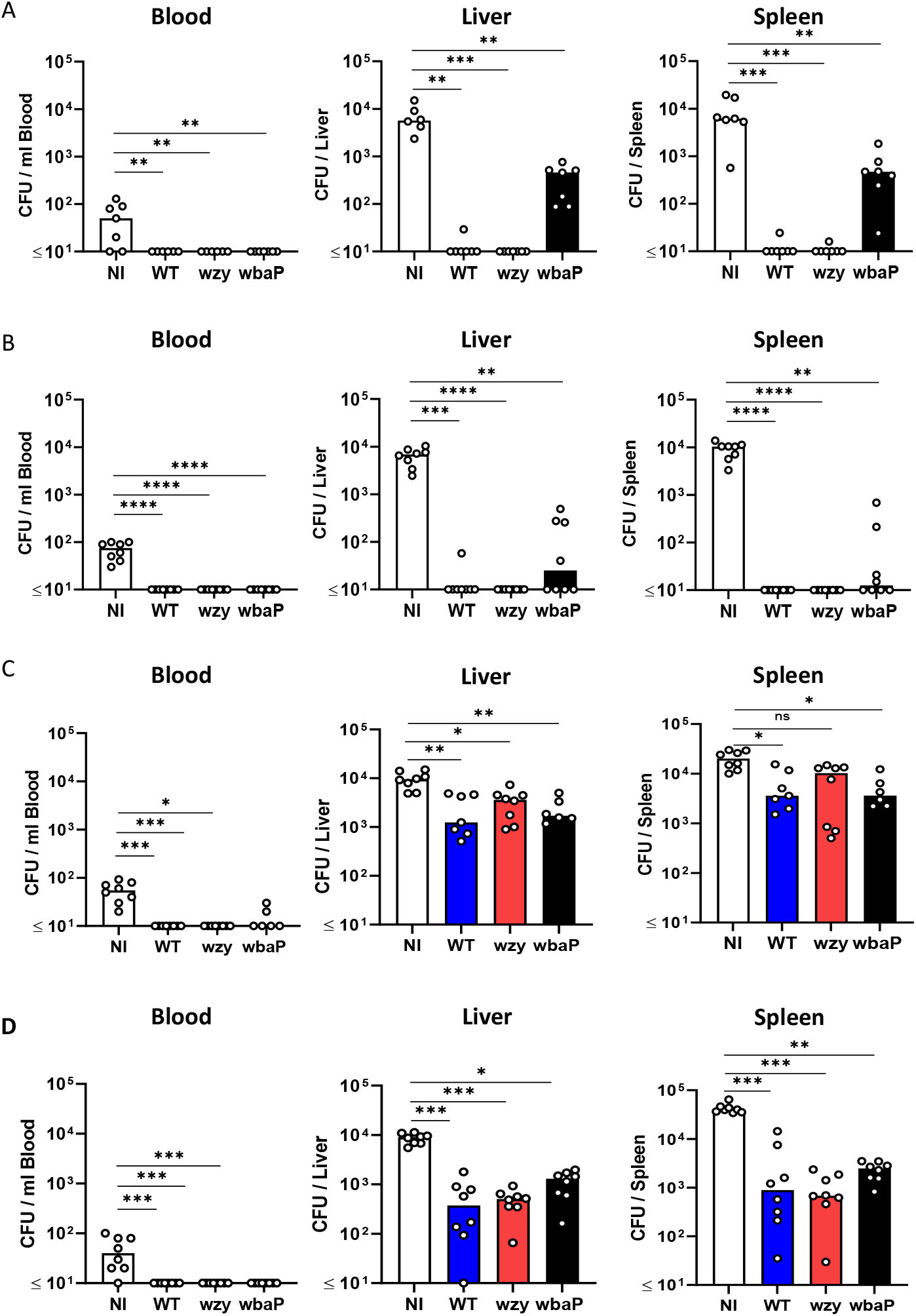
Bacterial colonization of tissues in immunized and challenged mice. wt mice were immunized i.p. with 1μg of OMV for 21 days or were boosted at 35 days for a further 21 days and challenged for 1 day with 10^4^ STm or SEn. **(A)** STm CFU in the blood (per mL), liver and spleen after primary immunization with OMV. **(B)** STm CFU in the blood (per mL), liver and spleen after secondary immunization with OMV. **(C)** SEn CFU in the blood (per mL), liver and spleen after primary immunization with OMV. **(D)** SEn CFU in the blood (per mL), liver and spleen after secondary immunization with OMV. Bars show medians and individual points represent results from a single mouse. NI = non-immunized, wt = wt-OMV, wzy = wzy-OMV, wbaP = wbaP-OMV. * = p≤0.05, ** = p≤0.01, *** = p≤0.005, **** = p≤0.001. Graphs show the results from 2 independent experiments. Elsner RA, Shlomchik MJ. Germinal Center and Extrafollicular B Cell Responses in Vaccination, Immunity, and Autoimmunity. *Immunity* **53**, 1136-1150 (2020).

## Discussion

Exposed antigens are the source of all licensed human bacterial vaccines. Understanding how antibodies bind surface antigens and contribute to controlling bacterial infections is essential for the development of novel vaccine strategies that reduce the burden of infectious disease and antimicrobial resistance. OMV are a clinically relevant vaccine platform and delivery technology that contains multiple antigens of value for such studies ^26^. We studied three OMV types differing in their LPS O-antigen content, as this antigen can, somewhat paradoxically, serve both as a target of protective antibodies and provide protection to the bacterium by occluding antibody access to other surface bacterial antigens ^7^. All OMV types induced similar total GC and IgG-expressing B cell responses, as well as ASC in the BM. Where the responses differed to each OMV, was in the distinct anti-O-antigen and anti-porin antibody signature.

The presence of O-antigen enhances the protection afforded by immunization with OMV and combined with its occluding role means there can be a hierarchy of protection afforded by antibodies to different surface antigens. For O-antigen, its surface exposure and presence at high density contributes to it being a key immunodominant antigen in non-capsular polysaccharide expressing Gram-negative bacteria. In contrast, other bacterial surface antigens may be immunodominant in the context of being well-recognised by antibodies after immunization or infection, but the antigen itself may be less accessible compared to O-antigen and thus may be less immunodominant in terms of protection. OmpA is an example of this^7^. Therefore, it is likely the combination of the level of antigen recognition by antibodies and the physical relationship between antigens that is critical for protection.

The lowest anti-LPS responses were typically observed after immunization with wbaP-OMV, consistent with the antibodies raised only being able to target core oligosaccharide and/or lipid A. In contrast, both the wt- and wzy-OMV induced similar serum IgG responses to LPS. A previous study had found that infection with live wzy-deficient *Salmonella* induced less anti-LPS antibody than the wt control ^30^. The differences observed in that study may reflect wzy-deficient *Salmonella* not persisting well in the host and so failing to provoke as robust a response as the wt bacteria. In contrast, using OMV as a vaccine platform overcomes issues associated with viability, such as concerns around using in immunocompromised populations, and help standardise the overall level of antigen encountered.

The reasons the two OMV types that contain different length O-antigens induce similar responses to LPS are unclear. It may be because a single O-antigen unit is sufficiently-sized to constitute an epitope ^31^, and thus longer O-antigen lengths are not necessary to generate equivalent levels of anti-LPS antibodies. This would mean that the number of LPS molecules within each OMV, rather than the length of the O-chain presented, drives the B-cell response. Alternatively, the strength of the anti-O-antigen response may simply reflect the intrinsic adjuvant properties of the OMV.

The control of STm infection was similar between the two O-antigen containing OMV groups, suggesting that the presentation of multiple O-antigen repeats is not essential for protection. Again, this may signify that a single O-antigen unit is sufficient to constitute a protective epitope or, alternatively, the protective epitope contains elements of both the O-antigen, and either the core oligosaccharide or an adjacent membrane antigen. This raises the key question of whether all O-antigen-containing epitopes are equivalent. Structural data and modelling studies suggest that the relationship between the O-antigen and the Fab itself is likely to be a critical factor in terms of determining the shape and size of the protective epitope ^31, 51, 49^. Notably, a number of structural studies reveal that epitopes derived from *Salmonella* and *Shigella* O-antigens comprise units as small as a trisaccharide ^31, 32, 51, 52^ supporting the view that antibodies to shorter, linear epitopes, induced by wzy-OMV, covering a single O-antigen unit, could plausibly provide protection to *Salmonella* expressing full-length O-antigen, as observed in this study.

Restricting O-antigen length did enhance antibody responses to non-O-antigen components of the OMV, illustrating the effectiveness of O-antigen at shielding the bacterial surface. Reduced bacterial burdens were observed after challenge of mice immunized with wbaP-OMV, indicating that non-O-antigen components of OMV induce responses can contribute to the restriction of bacterial numbers. Whilst it cannot be fully ruled out that these effects are due to antibodies targeting core oligosaccharide itself ^58^, the greater reduction in bacterial numbers after challenge with STm compared to SEn makes this less likely. Instead, we hypothesise that proteinaceous antigens within the OMV contribute to this. Some proteins that are known targets of such antibodies, e.g. OmpD ^17^, are present within the OMV, however additional antigens are likely to contribute to this effect.

Previous reports are not consistent in ascribing a protective value to levels of cross-reactive antibodies induced after immunization with OMV, OMPs, or bacteria expressing truncated LPS ^30, 59, 60, 61^. We found that boosting had a dramatic impact on levels of cross-protection, despite it only resulting in modest increases in anti-STm, LPS or porin-specific antibodies. Moreover, and counterintuitively, the greatest fold reduction in bacterial numbers was observed for the wt and wzy-OMV groups, rather than for the wbaP-OMV group. This would suggest that the improved efficacy is not due to changes in antibody titres against core oligosaccharide or lipid A, but rather due to the quality of the antibodies selected. Multiple reasons could explain these observations, none of which are mutually exclusive. One possibility is that boosting with wt or wzy-OMV increases antibodies against O-antigen species common to both STm and SEn (O1 and O12). The literature is limited on the value of such antibodies, but most studies suggest that anti-O12-antibodies are generally less protective than antibodies to the O4 and O9 moieties of STm and SEn respectively ^62 63 64^. Additionally, after natural infection there is little cross-protection observed ^6^, suggesting that anti-O1 or anti-O12-antibodies only make a minor contribution to protection. Otherwise, it is possible that O-antigen interacts with outer membrane surface proteins and helps maintain structural integrity, enhancing epitope stability and presentation. As boosting was the common factor in enhancing cross-protection, we hypothesise that secondary immunization increases cross-reactive antibodies with increased binding to conserved, but normally less accessible, epitopes through GC-mediated selection of B cells that do not target LPS core or O-antigen. This would rationalise the increased value of boosting, beyond simply increasing antibody levels.

In the context of invasive non-typhoidal *Salmonella* infections, the final formulation of an OMV vaccine will include OMV from both STm and SEn. Potentially, boosting with such an OMV vaccine can also enhance cross-protection to non-O4/9-expressing serovars. The proposed mechanism may also help explain the link between vaccination against group B meningococcus with vaccines that include OMV and the reduced risk of gonococcal infections ^65, 66^. Thus, OMV-based vaccines have great potential to be tailored to induce broadly-protective antibody responses against Gram-negative bacteria.

## Acknowledgements

This work was supported by grant C 82 1000664524 to AA and AFC. JCG is supported by National Institutes of Health grant R01-GM148586. Computational resources were provided through ACCESS (grant TG-MCB130173), which is supported by National Science Foundation grants 2138259, 2138286, 2138307, 2137603, and 2138296. Additional resources were provided by the Partnership for an Advanced Computing Environment (PACE) at the Georgia Institute of Technology.

## Author contributions

Performed experiments: A.A., S.E.J., M.P-T., E.D.M., R.P., A.E.M., D.N.L., G.G, J.P., F.E-R., K.A. Generated and provided novel reagents: J.A.B., M.B., C.L.M., I.R.H. Designed the experiments: D.N.L., I.R.H., J.C.G., J.Z.K., P.H., C.L.M., V.N.B., and A.F.C. Analysed the data: A.A., E.D.M., M.P-T., W.G.H., P.H., I.R.H., J.C.G., V.N.B., C.L.M., and A.F.C. Wrote the paper: A.A., M.P.T., J.Z.G., J.C.G., C.L.M., V.N.B., and A.F.C.

## Notes

### Competing Interest Statement

The authors have declared no competing interest.

