## Supplementary Materials for "Wild-type and single-O-antigen repeat outer-membrane vesicles induce equivalent protection against homologous and heterologous *Salmonella* challenge"

**Supplementary Table 1: Antibodies used for FACS.**

| Specificity and Fluorochrome | Stock Concentration | Dilution | Supplier | Staining panel/ Application |
|---|---|---|---|---|
| **Fcγ Receptor Blocking** | | | | |
| CD16:CD32 Purified | 0.5 mg/ml | 1:150 | eBioscience | Block FcγII and FcγIII receptors |
| **Extracellular Staining** | | | | |
| B220 Brilliant Violet 510 | 100 µg/ml | 1:200 | BioLegend | B cells Plasma cells |
| B220 APC eF-780 | 0.2 mg/ml | 1:300 | eBioscience | GC B cells |
| CD 19 Brilliant Violet 786 | 0.2 mg/ml | 1:400 | BD Biosciences | B cells GC B cells Plasma cells |
| CD138 Brilliant Violet 650 | 0.2 mg/ml | 1:200 | BD Biosciences | Plasma cells |
| CD38 Alexa Fluor 700 | 0.2 mg/ml | 1:200 | eBioscience | Plasma cells GC B cells panel |
| TACI Alexa Fluor 647 | 0.2 mg/ml | 1:200 | BD Biosciences | B cells panel |
| IgD Brilliant Violet 421 | 0.2 mg/ml | 1:150 | BioLegend | B cells panel |
| GL7 FITC | 0.6 mg/ml | 1:500 | BD Biosciences | GC B cells panel |
| FAS (CD95) Brilliant Violet 605 | 0.2 mg/ml | 1:100 | BD Biosciences | GC B cells panel |
| **Intracellular staining** | | | | |
| IgG FITC | 0.5 mg/ml | 1:500 | Southern Biotech | Plasma cells |
| IgM APC eF780 | 0.2 mg/ml | 1:300 | eBioscience | Plasma cells |
| BCL6 PE | 0.2 mg/ml | 1:500 | eBioscience | GC B cells panel |
| Ki-67 PE-Cy7 | 0.2 mg/ml | 1:200 | BD Biosciences | GC B cells panel |

**Supplementary Fig. 1. Protein profiles of wt, wzy, and wbaP-OMV.** OMV samples were digested with trypsin and mass spectrometric analysis performed. Proteins detected twice were recorded and listed based on their presence in one, two or all three of the OMV types. The distribution of these is shown in the diagram and individual proteins in the table.

**Supplementary Fig. 2. Flow cytometry gating strategy for GC B cells and intracellular IgM and IgG secreting cells. (A)** Singlets and then cells were isolated using forward and side scatter area and height parameters. B220 and CD19 were used to identify B cells. Germinal B cells (GCB) were identified as IgD- CD38+ GL-7+ Fas+ B cells. **(B)** Representative FACS plots for Fas+ GL7+ GC B cells in the spleen from immunized and non immunized mice. **(C)** Antibody secreting cells (ASC) were identified as TACI+ CD138+ single cells and separated as plasma cells (PC) and plasma blasts (PB) based on CD38 expression. IgG and IgM were used to identify the class of ASCs. **(D)** Representative FACS plots for Plasma cells stained for intracellular IgM and IgG, which were gated on plasma cells, in the spleen from immunized and non immunized mice.

**Supplementary Fig. 3. plasma(blast) cell proportions numbers induced after immunization with OMV. (A)** Proportions (left) and numbers (right) of intracellular $IgM^{hi}$ plasma(blast) cells in wt mice immunized i.p. with 1μg of OMV for 21 or 35 days or boosted at 35 days for a further 21 days. **(B)** Proportions (left) and numbers (right) of intracellular $IgG^{hi}$ plasma(blast) cells in the same mice. Bars show medians and individual points represent results from a single mouse. NI = non-immunized, wt = wt-OMV, wzy = wzy-OMV, wbaP = wbaP-OMV. * = p≤0.05, ** = p≤0.01, *** = p≤0.005, **** = p≤0.001. Graphs show the results from 2 independent experiments.

**Supplementary Fig. 4. Anti-LPS and anti-porin IgM antibody secreting cells in the spleen and Bone marrow induced to OMV.** wt mice were immunized i.p. with 1μg of OMV for 21 or 35 days or boosted at 35 days for a further 21 days. **(A)** Representative ELISPOT wells of the assessed spleen for anti-LPS and anti-porins IgM ASC (top), and anti-LPS and anti-porins IgG ASC (bottom) **(B)** Frequencies of splenic LPS or porin-specific IgM antibody-secreting cells determined by ELISPOT. **(C)** Frequencies of bone marrow LPS or porin-specific IgM antibody-secreting cells. Individual points represent spot forming units (SFU) from a single mouse. NI = non-immunized, wt = wt-OMV, wzy = wzy-OMV, wbaP = wbaP-OMV. * = p≤0.05, ** = p≤0.01, *** = p≤0.005. Graphs show the combined data from 2 independent experiments.

**wt-OMV – 2 hits** **wbaP-OMV – 2 hits**

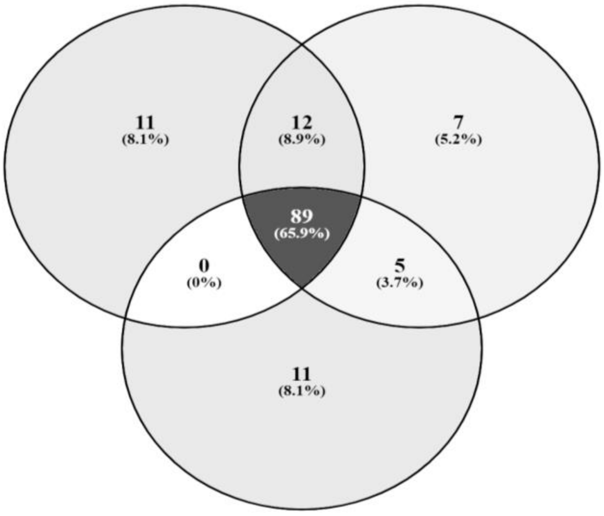


**wzy-OMV – 2 hits**

| wt only | Description |
|---|---|
| P52616 | FLJB_SALTY Phase 2 flagellin OS=Salmonella typhimurium (strain LT2 / SGSC1412... |
| P52615 | FLJB_SALAE Phase 2 flagellin OS=Salmonella abortus-equi OX=607 GN=fljB... |
| A9MFR9 | CH60_SALAR Chaperonin GroEL OS=Salmonella arizonae (strain ATCC BAA-731... |
| P0A1J1 | FLGE_SALTY Flagellar hook protein FlgE OS=Salmonella typhimurium (strain... |
| A9N0I9 | RL1_SALPB 50S ribosomal protein L1 OS=Salmonella paratyphi B (strain ATCC... |
| P35672 | SCTC1_SALTY SPI-1 type 3 secretion system secretin OS=Salmonella typhimurium... |
| P0A283 | PTGA_SALTY PTS system glucose-specific EIIA component OS=Salmonella typhimurium... |
| Q7CQY4 | CHIP_SALTY Chitoporin OS=Salmonella typhimurium (strain LT2 / SGSC1412... |
| A9MN68 | RL15_SALAR 50S ribosomal protein L15 OS=Salmonella arizonae (strain ATCC... |
| P0A297 | RL10_SALTY 50S ribosomal protein L10 OS=Salmonella typhimurium (strain... |
| Q8Z418 | PTRA_SALTI Protease 3 OS=Salmonella typhi OX=90370 GN=ptrA PE=3 SV=1 |

| wbap only | Description |
|---|---|
| Q56111 | OMPS2_SALTI Outer membrane protein S2 OS=Salmonella typhi OX=90370 GN=ompS2... |
| A9N8B8 | RL17_SALPB 50S ribosomal protein L17 OS=Salmonella paratyphi B (strain... |
| A9MPV4 | RS21_SALAR 30S ribosomal protein S21 OS=Salmonella arizonae (strain ATCC... |
| Q8Z740 | YNCE_SALTI Uncharacterized protein YncE OS=Salmonella typhi OX=90370 GN=yncE... |
| A9MN51 | RL2_SALAR 50S ribosomal protein L2 OS=Salmonella arizonae (strain ATCC... |
| A9MFK9 | RS18_SALAR 30S ribosomal protein S18 OS=Salmonella arizonae (strain ATCC... |
| P0A2A5 | RL28_SALTY 50S ribosomal protein L28 OS=Salmonella typhimurium (strain... |

| wzy only | Description |
|---|---|
| Q8ZP20 | TREA_SALTY Periplasmic trehalase OS=Salmonella typhimurium (strain LT2... |
| B5FTN7 | TREA_SALDC Periplasmic trehalase OS=Salmonella dublin (strain CT_02021853)... |
| Q7CPK0 | UGPB_SALTY sn-glycerol-3-phosphate-binding periplasmic protein UgpB OS=Salmonella... |
| Q8ZPD6 | YNCE_SALTY Uncharacterized protein YncE OS=Salmonella typhimurium (strain... |
| P06175 | FLIC_SALRU Flagellin OS=Salmonella rubislaw OX=598 GN=fliC PE=3 SV=2 |
| Q8Z289 | GUN_SALTI Endoglucanase OS=Salmonella typhi OX=90370 GN=bcsZ PE=3 SV=1 |
| D0ZSY7 | RSEB_SALT1 Sigma-E factor regulatory protein RseB OS=Salmonella typhimurium... |
| Q7CR85 | THIB_SALTY Thiamine-binding periplasmic protein OS=Salmonella typhimurium... |
| Q8ZQC4 | KCY_SALTY Cytidylate kinase OS=Salmonella typhimurium (strain LT2 / SGSC1412... |
| A9N7A9 | OPGB_SALPB Phosphoglycerol transferase I OS=Salmonella paratyphi B (strain... |
| P67714 | NDPA_SALTY Nucleoid-associated protein YejK OS=Salmonella typhimurium (strain... |

| wt + wbap | Description |
|---|---|
| P06179 | FLIC_SALTY Flagellin OS=Salmonella typhimurium (strain LT2 / SGSC1412 /... |
| A9MVX4 | TREA_SALPB Periplasmic trehalase OS=Salmonella paratyphi B (strain ATCC... |
| Q8ZNQ9 | WZA_SALTY Putative polysaccharide export protein wza OS=Salmonella typhimurium... |
| A9MRX7 | OPGB_SALAR Phosphoglycerol transferase I OS=Salmonella arizonae (strain... |
| A9MZ51 | RS16_SALPB 30S ribosomal protein S16 OS=Salmonella paratyphi B (strain... |
| A9N0R7 | RS2_SALPB 30S ribosomal protein S2 OS=Salmonella paratyphi B (strain ATCC... |
| A9MN53 | RL22_SALAR 50S ribosomal protein L22 OS=Salmonella arizonae (strain ATCC... |
| A9MNY0 | RL13_SALAR 50S ribosomal protein L13 OS=Salmonella arizonae (strain ATCC... |
| A9MN70 | RS11_SALAR 30S ribosomal protein S11 OS=Salmonella arizonae (strain ATCC... |
| Q5PJK8 | UGPB_SALPA sn-glycerol-3-phosphate-binding periplasmic protein UgpB OS=Salmonella... |
| O54297 | RS4_SALTY 30S ribosomal protein S4 OS=Salmonella typhimurium (strain LT2... |
| P40810 | ILVD_SALTY Dihydroxy-acid dehydratase OS=Salmonella typhimurium (strain... |

| wbap + wzy | Description |
|---|---|
| A9N876 | MSRP_SALPB Protein-methionine-sulfoxide reductase catalytic subunit MsrP... |
| Q5PK07 | RS13_SALPA 30S ribosomal protein S13 OS=Salmonella paratyphi A (strain... |
| A9N240 | RL20_SALPB 50S ribosomal protein L20 OS=Salmonella paratyphi B (strain... |
| Q8ZMB5 | PTRA_SALTY Protease 3 OS=Salmonella typhimurium (strain LT2 / SGSC1412... |
| A9MH07 | YCEI_SALAR Protein YceI OS=Salmonella arizonae (strain ATCC BAA-731 / CDC346-86... |

| wt + wbap + wzy | Description |
|---|---|
| P02936 | OMPA_SALTY Outer membrane protein A OS=Salmonella typhimurium (strain LT2... |
| Q8ZRP0 | BAMA_SALTY Outer membrane protein assembly factor BamA OS=Salmonella typhimurium... |
| Q8Z9A3 | BAMA_SALTI Outer membrane protein assembly factor BamA OS=Salmonella typhi... |
| B5BC62 | TOLB_SALPK Tol-Pal system protein TolB OS=Salmonella paratyphi A (strain... |
| Q8ZRW0 | LPTD_SALTY LPS-assembly protein LptD OS=Salmonella typhimurium (strain... |
| H9L451 | BAMB_SALTY Outer membrane protein assembly factor BamB OS=Salmonella typhimurium... |
| P37409 | BTUB_SALTY Vitamin B12 transporter BtuB OS=Salmonella typhimurium (strain... |
| P26982 | DEGP_SALTY Periplasmic serine endoprotease DegP OS=Salmonella typhimurium... |
| Q54001 | TOLC_SALEN Outer membrane protein TolC OS=Salmonella enteritidis OX=149539... |
| Q8ZLB8 | BCSC_SALTY Cellulose synthase operon protein C OS=Salmonella typhimurium... |
| P0A261 | TSX_SALTY Nucleoside-specific channel-forming protein Tsx OS=Salmonella... |
| A0A0H3NJI9 | OMPC_SALTS Outer membrane porin C OS=Salmonella typhimurium (strain... |
| Q5PDE6 | SURA_SALPA Chaperone SurA OS=Salmonella paratyphi A (strain ATCC 9150 /... |
| A0A0H3NBQ0 | OMPD_SALTS Outer membrane porin OmpD OS=Salmonella typhimurium (strain... |
| P37600 | PHSA_SALTY Thiosulfate reductase molybdopterin-containing subunit PhsA... |
| P26466 | LAMB_SALTY Maltoporin OS=Salmonella typhimurium (strain LT2 / SGSC1412... |
| A9N5G2 | NAPA_SALPB Periplasmic nitrate reductase OS=Salmonella paratyphi B (strain... |
| P0A1X0 | SLYB_SALTY Outer membrane lipoprotein SlyB OS=Salmonella typhimurium (strain... |
| P26265 | CPDB_SALTY 2',3'-cyclic-nucleotide 2'-phosphodiesterase/3'-nucleotidase... |
| P43669 | PRC_SALTY Tail-specific protease OS=Salmonella typhimurium (strain LT2... |
| Q56078 | BGLX_SALTY Periplasmic beta-glucosidase OS=Salmonella typhimurium (strain... |
| P19576 | MALE_SALTY Maltose/maltodextrin-binding periplasmic protein OS=Salmonella... |
| P0A2C7 | POTD_SALTY Spermidine/putrescine-binding periplasmic protein OS=Salmonella... |
| Q8ZNA5 | FADL_SALTY Long-chain fatty acid transport protein OS=Salmonella typhimurium... |
| Q8ZP50 | OMPW_SALTY Outer membrane protein W OS=Salmonella typhimurium (strain LT2... |
| Q7CQN4 | LPP1_SALTY Major outer membrane lipoprotein Lpp 1 OS=Salmonella typhimurium... |
| P40827 | NLPD_SALTY Murein hydrolase activator NlpD OS=Salmonella typhimurium (strain... |
| Q8ZR01 | RLPA_SALTY Endolytic peptidoglycan transglycosylase RlpA OS=Salmonella... |
| B5R535 | OPGD_SALEP Glucans biosynthesis protein D OS=Salmonella enteritidis PT4... |
| P26478 | MALM_SALTY Maltose operon periplasmic protein OS=Salmonella typhimurium... |
| P0A2H9 | DSBA_SALTY Thiol:disulfide interchange protein DsbA OS=Salmonella typhimurium... |
| P06202 | OPPA_SALTY Periplasmic oligopeptide-binding protein OS=Salmonella typhimurium... |
| Q8ZR06 | PAGP_SALTY Lipid A palmitoyltransferase PagP OS=Salmonella typhimurium... |
| Q7CPZ3 | BAME_SALTY Outer membrane protein assembly factor BamE OS=Salmonella typhimurium... |
| P0A231 | PA1_SALTY Phospholipase A1 OS=Salmonella typhimurium (strain LT2 / SGSC1412... |
| P02910 | HISJ_SALTY Histidine-binding periplasmic protein OS=Salmonella typhimurium... |
| A9N4R0 | MLTC_SALPB Membrane-bound lytic murein transglycosylase C OS=Salmonella... |
| P39434 | SLT_SALTY Soluble lytic murein transglycosylase OS=Salmonella typhimurium... |
| Q57JI4 | NLPI_SALCH Lipoprotein NlpI OS=Salmonella choleraesuis (strain SC-B67)... |
| P50335 | YHJJ_SALTY Protein YhjJ OS=Salmonella typhimurium (strain LT2 / SGSC1412... |
| A0A0H3N9T8 | OMPF_SALTS Outer membrane porin F OS=Salmonella typhimurium (strain... |
| A9MZG4 | LSRB_SALPB Autoinducer 2-binding protein LsrB OS=Salmonella paratyphi B... |
| A9MVX1 | EMTA_SALPB Endo-type membrane-bound lytic murein transglycosylase A OS=Salmonella... |
| E1WAU4 | TAMA_SALTS Translocation and assembly module subunit TamA OS=Salmonella... |
| Q56030 | ENVE_SALTY Probable lipoprotein EnvE OS=Salmonella typhimurium (strain... |

| wt + wbap + wzy | Description |
|---|---|
| P37723 | OSMB_SALTY Osmotically-inducible lipoprotein B OS=Salmonella typhimurium... |
| P0A2C5 | RBSB_SALTY Ribose import binding protein RbsB OS=Salmonella typhimurium... |
| P22107 | TRAT_SALTM TraT complement resistance protein OS=Salmonella typhimurium... |
| Q06399 | YEDD_SALTY Uncharacterized lipoprotein YedD OS=Salmonella typhimurium (strain... |
| P06196 | USHA_SALTY Silent protein UshA(0) OS=Salmonella typhimurium (strain LT2... |
| Q8ZQZ7 | LPTE_SALTY LPS-assembly lipoprotein LptE OS=Salmonella typhimurium (strain... |
| P23905 | DGAL_SALTY D-galactose-binding periplasmic protein OS=Salmonella typhimurium... |
| P55890 | DSBC_SALTY Thiol:disulfide interchange protein DsbC OS=Salmonella typhimurium... |
| O33921 | AGP_SALTY Glucose-1-phosphatase OS=Salmonella typhimurium (strain LT2 /... |
| Q8ZQ08 | LPOB_SALTY Penicillin-binding protein activator LpoB OS=Salmonella typhimurium... |
| A9MP99 | LOLB_SALAR Outer-membrane lipoprotein LolB OS=Salmonella arizonae (strain... |
| P20753 | PPIA_SALTY Peptidyl-prolyl cis-trans isomerase A OS=Salmonella typhimurium... |
| A9N7X9 | LOLA_SALPB Outer-membrane lipoprotein carrier protein OS=Salmonella paratyphi... |
| Q8ZRJ9 | PAGN_SALTY Outer membrane protein PagN OS=Salmonella typhimurium (strain... |
| B5R236 | ECOT_SALEP Ecotin OS=Salmonella enteritidis PT4 (strain P125109) OX=550537... |
| P0A1I1 | PHSB_SALTY Thiosulfate reductase electron transfer subunit PhsB OS=Salmonella... |
| P58683 | APHA_SALTY Class B acid phosphatase OS=Salmonella typhimurium (strain LT2... |
| P23988 | PAGC_SALTY Virulence membrane protein PagC OS=Salmonella typhimurium (strain... |
| P64437 | YBGS_SALTY Uncharacterized protein YbgS OS=Salmonella typhimurium (strain... |
| P0CW86 | SODC1_SALTY Superoxide dismutase [Cu-Zn] 1 OS=Salmonella typhimurium (strain... |
| P0A1Z2 | SKP_SALTY Chaperone protein Skp OS=Salmonella typhimurium (strain LT2 /... |
| B5BBD9 | OPGG_SALPK Glucans biosynthesis protein G OS=Salmonella paratyphi A (strain... |
| P26976 | PHON_SALTY Non-specific acid phosphatase OS=Salmonella typhimurium (strain... |
| A9MUZ0 | YEBF_SALPB Protein YebF OS=Salmonella paratyphi B (strain ATCC BAA-1250... |
| P0A1C5 | FTSP_SALTY Cell division protein FtsP OS=Salmonella typhimurium (strain... |
| P0A1T6 | YIFL_SALTY Uncharacterized lipoprotein YifL OS=Salmonella typhimurium (strain... |
| A9MHG0 | EFTU_SALAR Elongation factor Tu OS=Salmonella arizonae (strain ATCC BAA-731... |
| P66950 | BEPA_SALTY Beta-barrel assembly-enhancing protease OS=Salmonella typhimurium... |
| P40680 | ERFK_SALTY Probable L,D-transpeptidase ErfK/SrfK OS=Salmonella typhimurium... |
| O68901 | SODC_SALTY Superoxide dismutase [Cu-Zn] 2 OS=Salmonella typhimurium (strain... |
| P63727 | C562_SALTY Soluble cytochrome b562 OS=Salmonella typhimurium (strain LT2... |
| Q57I62 | YICS_SALCH Uncharacterized protein YicS OS=Salmonella choleraesuis (strain... |
| Q8ZML1 | PROX_SALTY Glycine betaine/proline betaine-binding periplasmic protein... |
| A9MR48 | RS20_SALAR 30S ribosomal protein S20 OS=Salmonella arizonae (strain ATCC... |
| A9MRN3 | YNFB_SALAR UPF0482 protein YnfB OS=Salmonella arizonae (strain ATCC BAA-731... |
| Q8ZQX4 | CHIQ_SALTY Uncharacterized lipoprotein ChiQ OS=Salmonella typhimurium (strain... |
| Q8ZQM3 | GSIB_SALTY Glutathione-binding protein GsiB OS=Salmonella typhimurium (strain... |
| P26366 | AMIB_SALTY N-acetylmuramoyl-L-alanine amidase AmiB OS=Salmonella typhimurium... |
| Q9ZF60 | GLTI_SALTY Glutamate/aspartate import solute-binding protein OS=Salmonella... |
| P0CL43 | SCTG_SALTY SPI-1 type 3 secretion system pilotin OS=Salmonella typhimurium... |
| Q8ZRS2 | CUEO_SALTY Multicopper oxidase CueO OS=Salmonella typhimurium (strain LT2... |
| A9MGK5 | NRFA_SALAR Cytochrome c-552 OS=Salmonella arizonae (strain ATCC BAA-731... |
| A9MN54 | RS3_SALAR 30S ribosomal protein S3 OS=Salmonella arizonae (strain ATCC... |
| P66643 | RS9_SALTY 30S ribosomal protein S9 OS=Salmonella typhimurium (strain LT2... |

Supplementary Fig. 1.

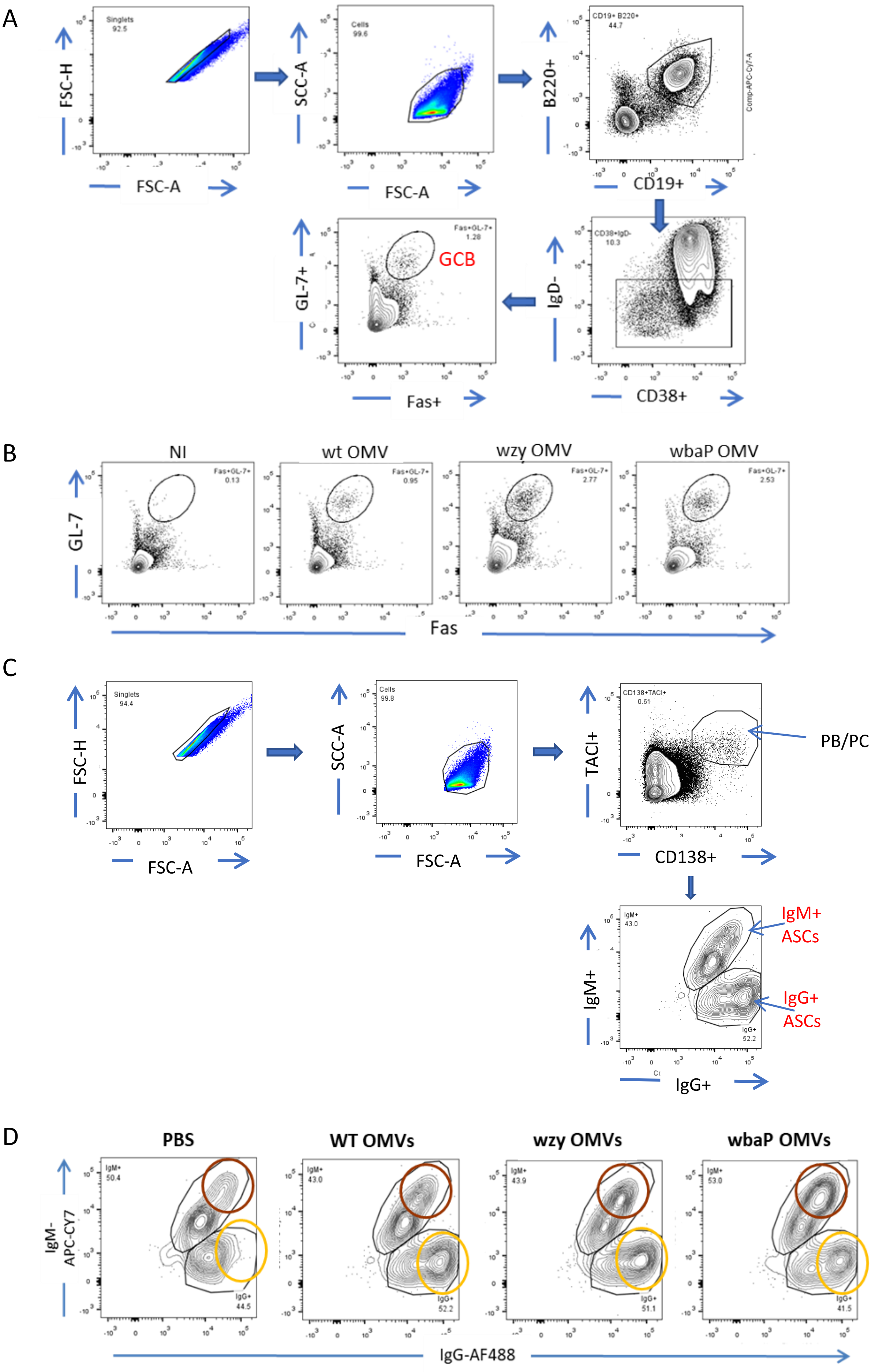


Supplementary Fig. 2.

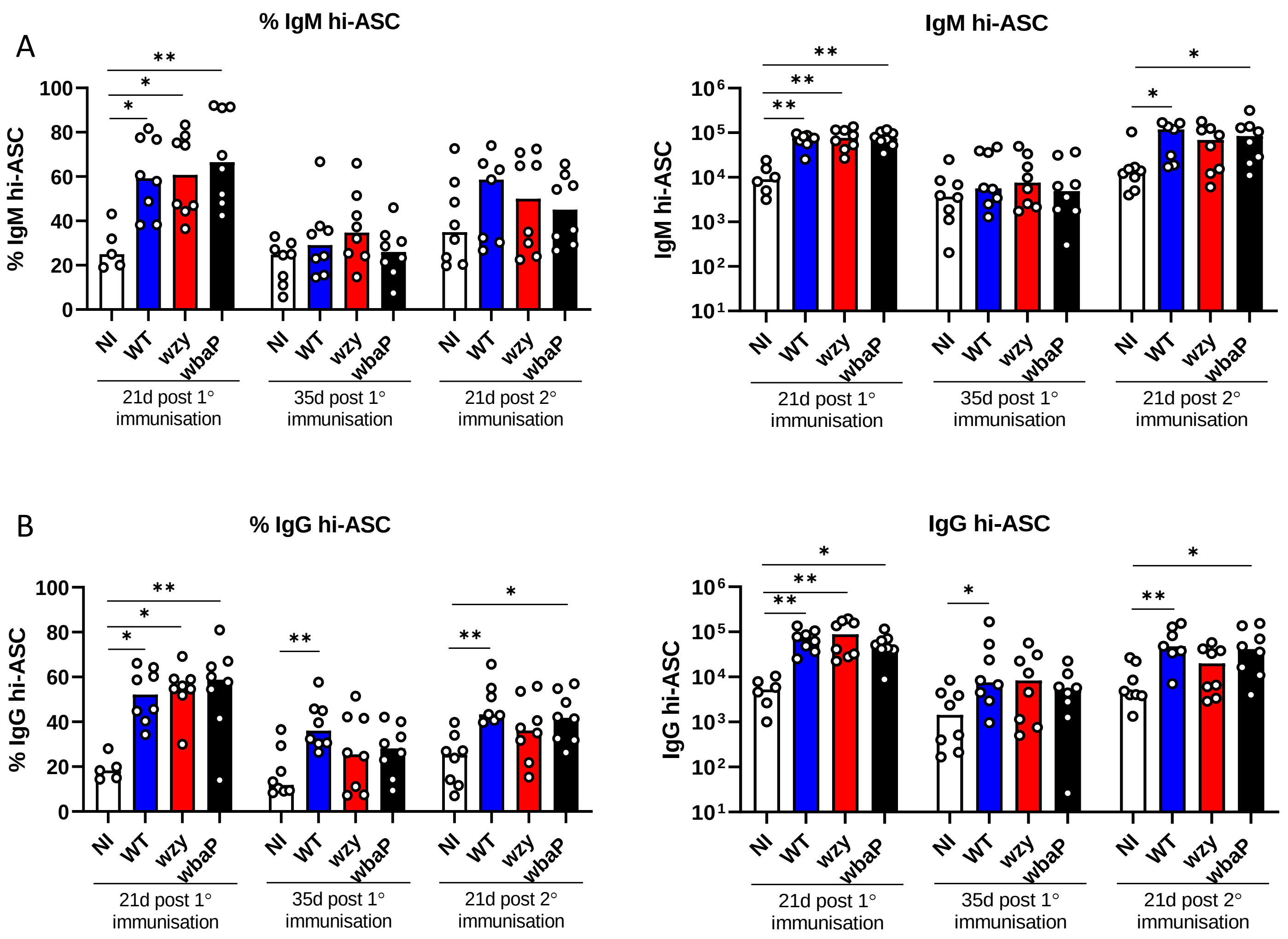


Supplementary Fig. 3.

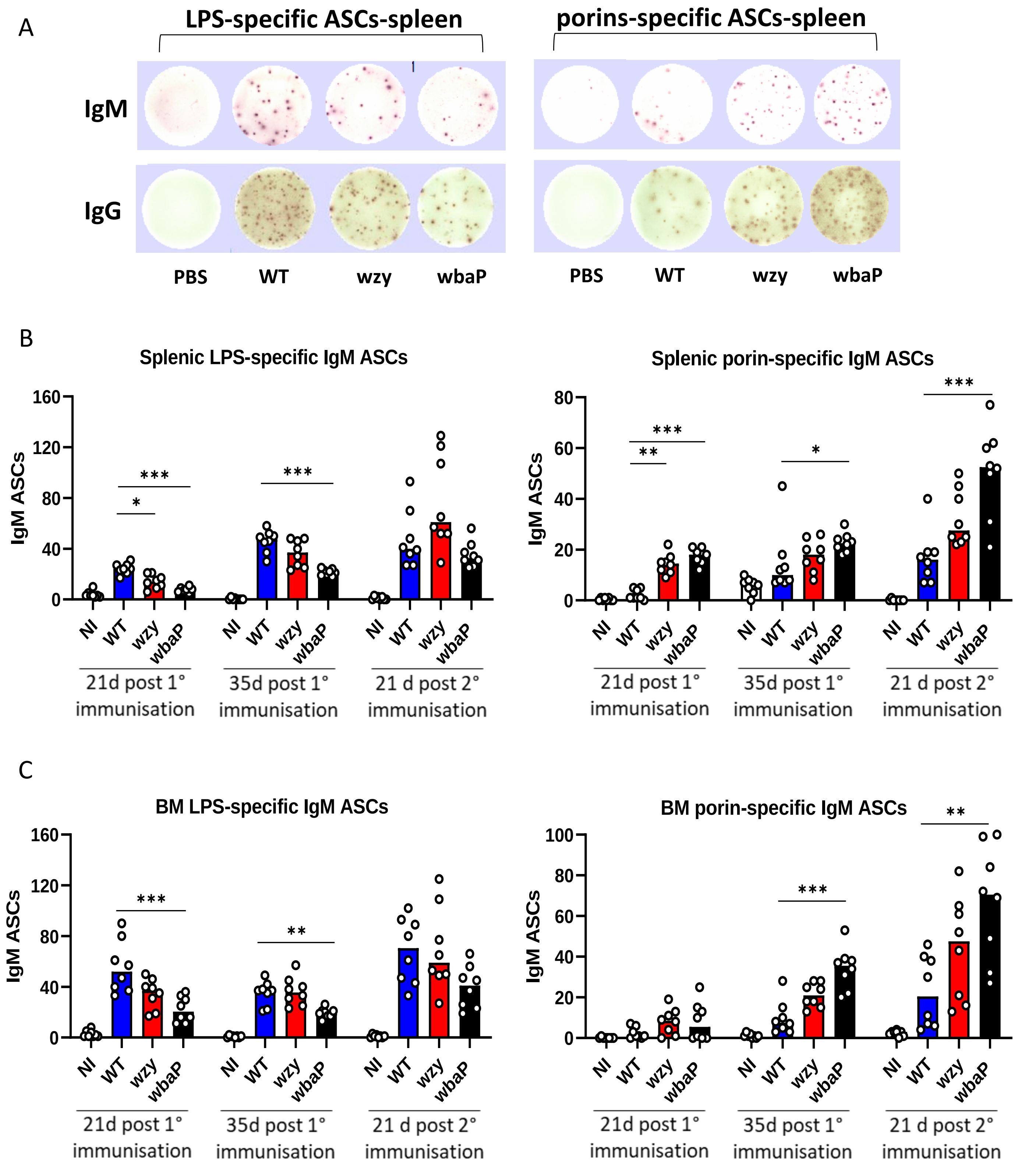


Supplementary Fig. 4.
